# Loss of DNA Demethylase Enhances Arsenic Resistance Via Reducing of *PIN2* Antisense Long Noncoding RNA

**DOI:** 10.64898/2026.09.25.754583

**Authors:** Yuxin Wang, Xinyan Zhou, Zhengjie Jiang, Zijun Xia, Zishan Huang, Litao Wang, Qi Mi, Yuxin Hou, Yibo Wang, Keqin Miao, Yi’an Chen, Hanchen Ye, Jingyuan Zhang, Haoyuan Gu, Luning Zeng, Hengyu Shen, Shang Wang, Yang Yu, Junran Ye, Ruoxuan Wang, Yuan Yao, Yitong Wang, Joohyun Lee

## Abstract

DNA methylation dynamics regulate diverse plant stress responses, but the mechanisms governing arsenic (As) tolerance through epigenetic pathways remain largely unknown. Here we report that Arabidopsis mutants lacking the DNA demethylase, REPRESSOR OF SILENCING (ROS1), show substantially enhanced As resistance. Integrating transcriptome and methylome data, we found that *PIN-FORMED 2* (*PIN2*), encoding an auxin efflux carrier that also transports As out of cells, represents a key ROS1 target. In *ros1*, we observed increased methylation in the last exon of *PIN2*, a region functioning as the promoter for an antisense long non-coding RNA (lncRNA). Hypermethylation of the promoter suppresses the antisense lncRNA, which follows reduction of sense *PIN2* transcription. As a result, *PIN2* expression increases in *ros1* mutants during As exposure, enhancing As tolerance. Analysis of *ros1 pin2* double mutants confirmed that PIN2 acts downstream of ROS1 in As tolerance. Our study demonstrates an epigenetic pathway in which ROS1 regulates As tolerance by controlling *PIN2* expression through antisense lncRNA-mediated transcriptional interference, offering new epigenetic strategies using plants in heavy metal contaminated environments.

## Main Text

Arsenic (As) is a toxic metalloid that threatens human health and crop production worldwide (*1–3*). Elemental As has low solubility and toxicity, but its oxidized forms, arsenate [As(V)] and arsenite [As(III)], dissolve easily and harm living organisms (*4–6*). Both natural processes and human activities release As into the environment. Weathering of As-bearing rocks, mining, farming, and groundwater use mobilize As from soils into water (*7–9*), as well as industrial waste from alloy, electronics, and glass production (*8*). Poor regulation of groundwater and waste has caused widespread As contamination in soils and water globally (*10*). In the United States, As levels in drinking water have dropped slightly since 2006, but many small groundwater systems in 25 states still exceed the EPA limit of 10 µg/L (*11–13*). The problem is worse in Asia, where hundreds of millions in Bangladesh, India, and China face high As exposure through contaminated drinking water and irrigated crops (*14, 15*). Chronic As exposure causes skin lesions, digestive disorders, heart and lung diseases, and cancers, leading the Centers for Disease Control and Prevention (CDC) to classify As as a carcinogen (*16*).

Plants are among the first organisms exposed to As in contaminated environments and serve as entry points into food chains (*17, 18*). Since As(V) resembles phosphate chemically, plants take it up through phosphate transporters and reduce it to the more toxic As(III) inside cells (*19, 20*). As disrupts enzymes by replacing essential metal cofactors and generates harmful reactive oxygen species (ROS) (*5, 21, 22*), suppressing plant growth, lowering crop yields, and damaging vegetation important for soil and water conservation (*23, 24*). To survive As stress, plants use transport control and chelation. Aquaporins like OsNIP (NODULIN26- LIKE INTRINSIC PROTEIN) in rice and AtPIP (PLASMA MEMBRANE INTRINSIC PROTEIN) in Arabidopsis regulate As uptake, while transporters such as AtPIN2 (PIN-FORMED 2) export As from cells (*24–27*). For detoxification, plants synthesize sulfur-rich molecules—glutathione (GSH) and phytochelatins (PCs)—that chelate As tightly (*28*). In Arabidopsis, AtGS1 and AtGS2 (GLUTATHIONE SYNTHASE 1 and 2) produce GSH, while AtPCS1 and AtPCS2 (PHYTOCHELATIN SYNTHASE 1 and 2) make PCs (*29, 30*).

These chelators capture As and sequester it into vacuoles, reducing cytoplasmic toxicity (*31–33*).

Plant responses to As and other heavy metals depend on both genetic and epigenetic controls (*34, 35*). DNA methylation is a key epigenetic mechanism that regulates gene expression, with effects depending on where methylation occurs and the sequence context (*36, 37*).

Despite progress in understanding DNA demethylases in heavy metal stress responses, no direct target genes regulated by DNA demethylation have been confirmed experimentally (*38–40*). To address this question, we screened Arabidopsis DNA methylation mutants and identified that loss of *ROS1* (*REPRESSOR OF SILENCING 1*), which encodes a DNA demethylase, confers As tolerance. We combined transcriptome and methylome analyses to identify *ROS1*-regulated genes affecting As tolerance. We found that loss of *ROS1* increases both *PIN2* expression and DNA methylation at the 3′ untranslated region (3′ UTR) of the *PIN2* gene, improving As tolerance. Genetic analysis using *pin2 ros1* double mutants confirmed that PIN2 acts downstream of ROS1 in As tolerance. Notably, the methylation changes at the *PIN2* 3′ UTR—which serves as a promoter for an antisense long noncoding RNA (lncRNA)—reducing this antisense transcript that normally represses *PIN2* expression. Specifically, hypermethylation of the antisense lncRNA promoter in *ros1* mutants blocks antisense RNA production, leading to increased *PIN2* expression and enhanced As tolerance. Our results reveal an epigenetic pathway in which ROS1-mediated DNA demethylation regulates plant As tolerance by controlling *PIN2* through an antisense lncRNA, providing new insights for improving plants on contaminated land.

## Defective DNA demethylation machinery elevates As tolerance in Arabidopsis

In Arabidopsis, active DNA demethylation depends mainly on a family of DNA glycosylases: ROS1, DEMETER-LIKE2 (DML2), and DEMETER-LIKE3 (DML3) (*41, 42*). Earlier work linked DNA demethylation to cadmium (Cd) tolerance, showing that the triple mutant lacking all three demethylases displayed enhanced Cd resistance in Arabidopsis (*40*). However, those studies did not identify which demethylase was responsible or pinpoint the downstream targets where increased methylation drives tolerance. We therefore investigated whether DNA demethylation also affects As stress responses in Arabidopsis.

We started by screening single mutants for each demethylase under As treatment. Among these, only *ros1-4* showed clear resistance to As compared to wild-type plants (Fig. 1A-C).

**Fig. 1.**
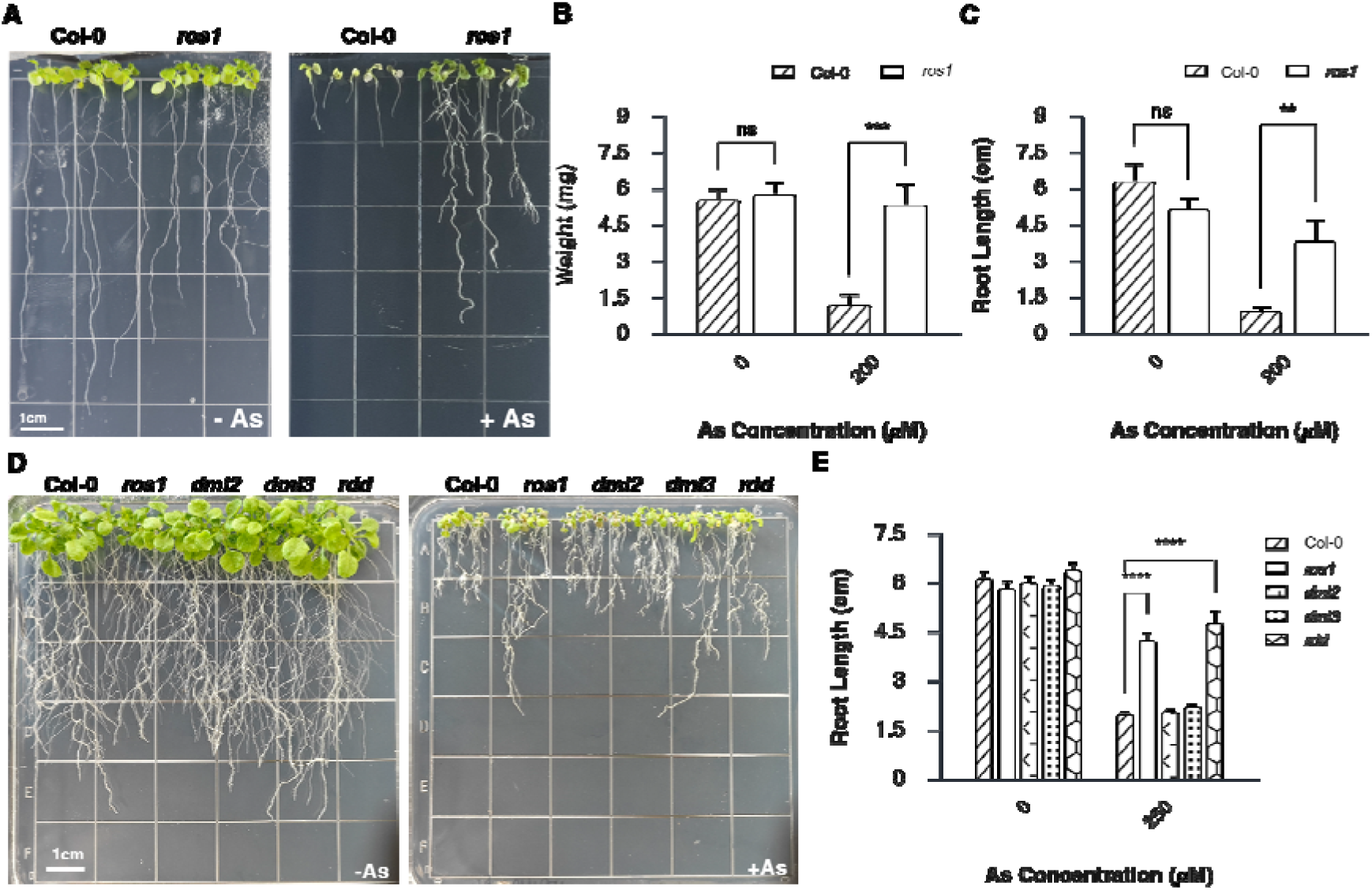
As tolerance in Arabidopsis DNA demethylase mutants. (**A**) Growth phenotypes of wild-type and *ros1* plants grown on half-strength MS medium without (left) or with (right) 200 μM arsenic. Scale bar = 1 cm. (**B**) Fresh weight and (**C**) primary root length of seedlings shown in (A). (**D**) Growth phenotypes of wild-type, *ros1*, *dml2*, *dml3*, and *rdd* mutants under control (left) and As stress (right) conditions. Scale bar = 1 cm. (**E**) Root length comparison among wild-type, *ros1*, *dml2*, *dml3*, and *rdd* mutants with or without 250 μM As. Data were analyzed by two-way ANOVA. Asterisks denote statistical significance: \*\*\*\**P* < 0.0001, \*\*\**P* < 0.001, \*\**P* < 0.01; ns, not significant.

To test whether DML2 or DML3 might act redundantly with ROS1, we examined *dml2-2* and *dml3-2* single mutants as well as the *ros1-4 dml2-2 dml3-2* triple mutant (*rdd*) (Fig. 1D). Quantitative measurements revealed that *ros1* and *rdd* mutants exhibited statistically similar levels of As tolerance, both significantly higher than wild type (Fig. 1E). In contrast, *dml2* and *dml3* single mutants remained as sensitive to As as wild-type plants. These findings demonstrate that ROS1 serves as the principal demethylase governing As regulation in Arabidopsis, while DML2 and DML3 contribute minimally to As stress response. The observation that *rdd* triple mutants displayed As tolerance comparable to *ros1* single mutants further supports ROS1’s dominant role, suggesting limited functional redundancy among demethylase family members under As stress conditions.

## ROS1-mediated demethylation directly regulates As tolerance independent of transposon mobilization

To investigate how the loss of *ROS1* contributes to As tolerance, we performed whole transcriptome sequencing (RNA-seq) and whole genome bisulfite sequencing (WGBS) using wild-type, *ros1* and *rdd* with and without As treatment to assess global and locus-specific changes in gene expression and DNA methylation levels. In addition, to identify the sole effect of ROS1, we also added *dml2*, *dml3* and *rdd* to narrow down to the key targets contributing to As tolerance in these analyses.

In *ros1* and *rdd*, which lack functional *ROS1*, the global transcriptomic response to As treatment were similarly altered compared to those in wild type, *dml2*, and *dml3* (Fig. 2A). Compared to wild type under As treatment, *ros1* exhibited 360 up-regulated and 196 down- regulated differentially expressed genes (DEGs), and *rdd* showed 987 up-regulated and 208 down-regulated DEGs, highlighting extensive transcriptomic alterations in both mutants (Fig. 2B). In contrast, *dml2* and *dml3* exhibited substantially fewer transcriptional changes under As treatment, exhibiting limited transcriptional changes of these demethylases in the As response (Fig. 2B and fig. S1). By comparison, *ros1* and *rdd* showed similar transcriptomic alterations under As treatment, with 337 DEGs shared between only these two mutants, while the overlap with *dml2* and *dml3* is minimal (Fig. 2C).

**Fig. 2.**
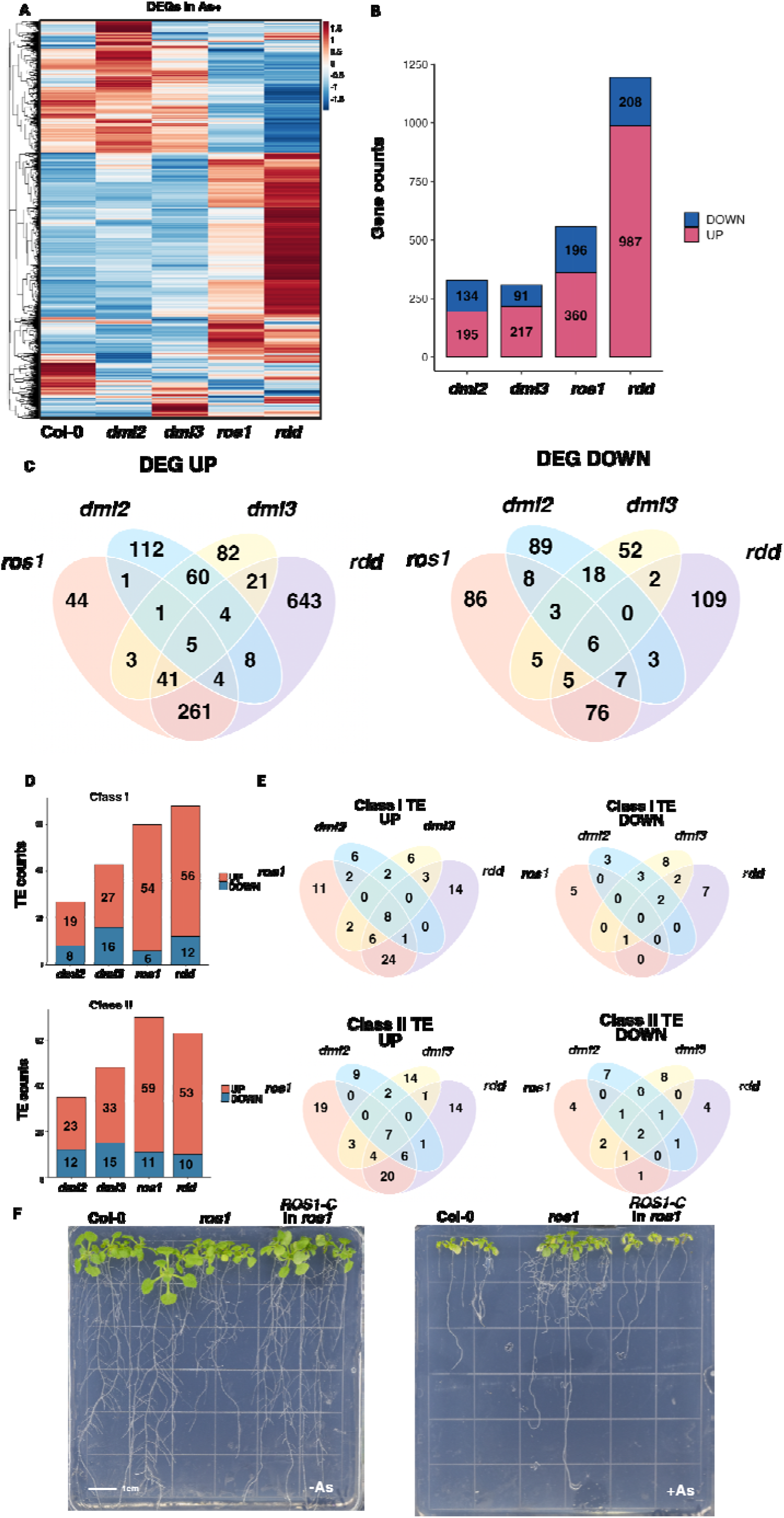
Transcriptional profiling of DNA demethylase mutants under As stress and *ROS1* complementation in *ros1*. (**A**) DEGs in *dml2*, *dml3*, *ros1*, and *rdd* mutants versus wild-type after exposure to 250 μM As. (**B**) Total numbers of upregulated and downregulated DEGs in *dml2*, *dml3*, *ros1*, and *rdd* mutants. (**C**) Venn diagram illustrating DEG overlap among the four mutant lines. (**D**) Total numbers of differentially expressed Class I and Class II TEs showing upregulation or downregulation in each mutant. (**E**) Distribution of differentially expressed Class I and Class II TEs among DNA demethylase mutants. (**F**) Growth phenotypes of wild-type (Col-0), *ros1*, and complemented *ros1* (*ROS1-C*) under control conditions (left) and As treatment (right).

To further investigate the relevance of genes differentially expressed in response to As stress, we performed gene ontology (GO) analysis and gene set enrichment analysis (GSEA) on DEGs identified across all demethylase mutants. In both *ros1* and *rdd*, DEGs were correlated including abiotic stress responses (Fig. S2 and Table S2). These enrichment patterns correspond with the distinct transcriptomic and phenotypic profiles of the mutants, indicating that *ROS1*-deficient lines, *ros1* and *rdd*, display similar transcriptional responses to As stress compared with *dml2* and *dml3*.

Loss of *ROS1* causes widespread DNA methylation can affect both gene expression and transposable element (TE) activity. TEs are mobile sequences classified as Class I transposons (retrotransposons) that mobilize via an RNA intermediate, or Class II transposons (DNA transposons) that move by a cut-and-paste mechanism (*43, 44*). Since these elements can insert randomly throughout the genome, we considered whether TE mobilization in *ros1* mutants might create As tolerance by disrupting relevant genes. We therefore monitored Class I and Class II transposon expression and observed predominantly increased expression in DNA demethylase mutants after As treatment (Fig. 2, D and E, and Fig. S3).

Reactivated transposons could contribute to As tolerance through genomic insertions that alter gene expression in a *cis* or *trans* manner via promoter interference, transcriptional readthrough, or chromatin remodeling (*45–48*). To distinguish whether As tolerance in *ros1* is caused by direct transcriptional changes or by transposon mobilization, we performed complementation experiments by reintroducing functional *ROS1* into the *ros1* mutant that was tested under As in Fig 1A. Transposon-induced changes involve stable genomic insertions that persist independently of ROS1 activity, whereas methylation-dependent regulation should reverse upon ROS1 restoration. The complemented lines displayed As sensitivity comparable to wild type, indicating that enhanced tolerance results from reversible methylation-dependent gene expression rather than heritable transposon-induced alterations (Fig. 2F). These data demonstrate that ROS1-mediated demethylation directly controls As stress responses through targeted regulation of specific genes.

## ROS1 preferentially targets mainly CG methylation in gene regions under As Stress

To determine whether the transcriptional changes in *ros1* mutants result from altered DNA methylation patterns, we performed whole-genome bisulfite sequencing (WGBS) on DNA demethylase mutants and wild-type plants following As exposure. The methylation profiles in *ros1* and *rdd* mutants aligned with their distinct gene expression patterns and As-resistant phenotypes (Fig. 3A, and Fig. S4 and S5). Quantitative analysis revealed that CG methylation accounted for the majority of changes, with approximately 73.5% of total methylation alterations in *ros1* plants under As treatment, whereas CHG and CHH methylation represented only 18.6% and 7.9%, respectively (Fig. 3B).

**Fig. 3.**
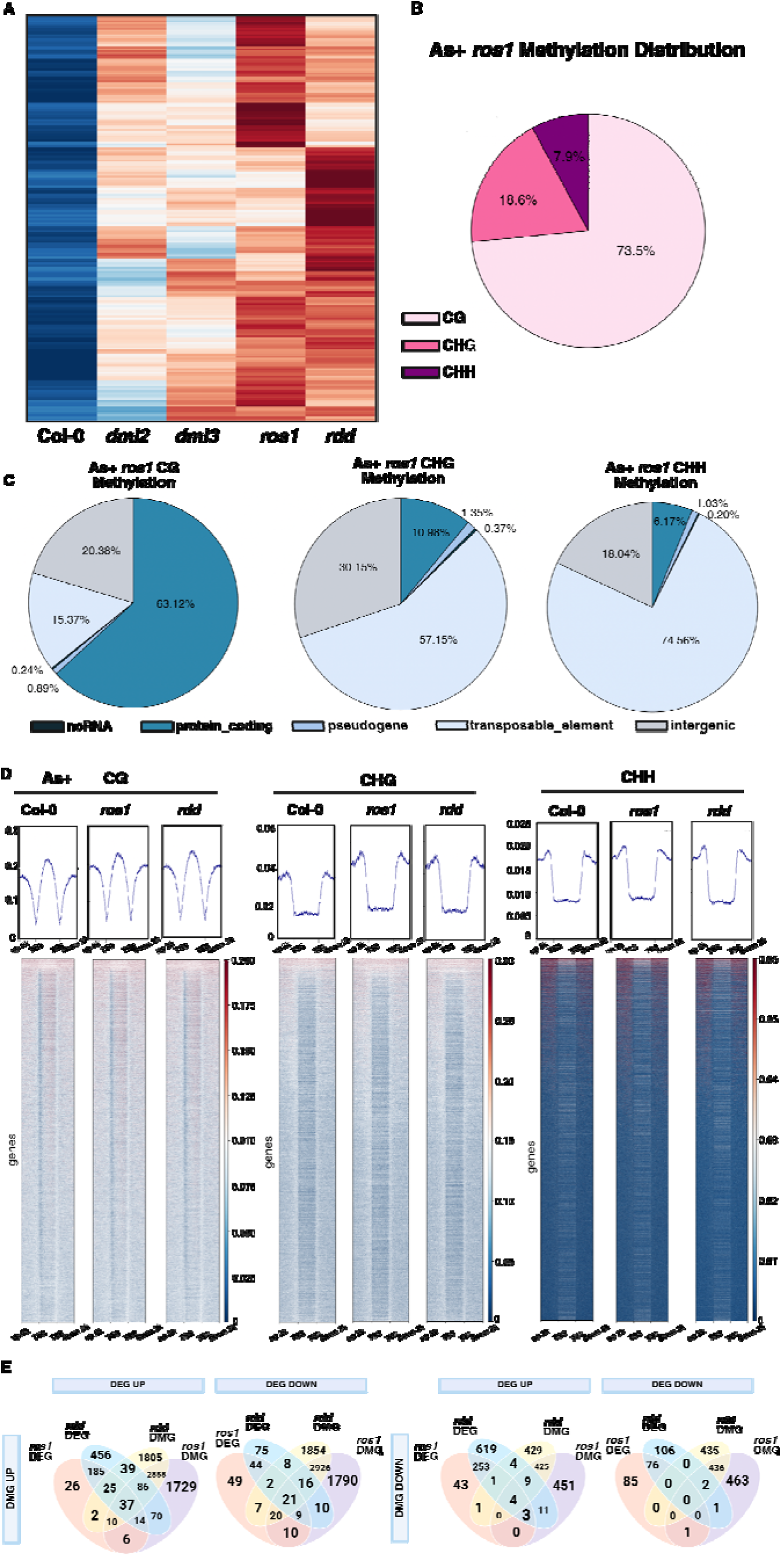
Methylomic profiling of DNA demethylase mutants under As stress and integration with transcriptome. (**A**) DMGs identified in *dml2*, *dml3*, *ros1*, and *rdd* mutants relative to wild type under As stress. (**B**) Distribution of methylation types (CG, CHG, and CHH) among DMGs in *ros1* under As stress. (**C**) Distribution of DMGs for CG, CHG and CHH contexts across different genomic regions in *ros1*. (**D**) Methylation densities for CG, CHG, and CHH contexts spanning gene bodies and flanking 2 kb sequences in wild-type, *ros1*, and *rdd* under As stress. TSS, transcription start site; TES, transcription end site. (**E**) Integration of expression data (DEG UP, upregulated; DEG DOWN, downregulated) with methylation data (DMG UP, hypermethylated; DMG DOWN, hypomethylated) in *ros1* and *rdd*.

The genomic distribution of these methylation changes showed that CG methylation changes concentrated heavily in gene bodies and promoter regions (63.12%), while CHG and CHH methylation changes in these regions were substantially lower (10.98% and 6.17%, respectively) (Fig. 3C). Within gene regions, DNA methylation differences in *ros1* mutants extended across promoters, transcription start sites, gene bodies, and downstream regions (Fig. 3D). Both *ros1* and *rdd* mutants displayed CG and CHG methylation alterations throughout these genomic features, suggesting comprehensive regulation of gene-associated methylation. The parallel increases in methylation and corresponding gene expression changes in *ros1* and *rdd* mutants strongly suggest that *ROS1* directly regulates As-responsive genes through demethylation. These findings establish that *ROS1* primarily removes mainly CG methylation from gene-associated regions, including both promoters and gene bodies, which proves essential for appropriate transcriptional responses during As stress.

## *PIN2* is the primary downstream target of ROS1 in As tolerance

To identify genes whose expression depends on *ROS1*-mediated demethylation during As stress, we systematically compared differentially expressed genes (DEGs) with differentially methylated genes (DMGs) in *ros1* and *rdd* mutants following As treatment, excluding *dml2* and *dml3* to focus on *ROS1*-specific effects. Although various genes showed either expression or methylation changes, relatively few displayed coordinated alterations in both parameters (Fig. 3E). The majority of overlapping genes were both upregulated and hypermethylated—we detected 37 such genes only in both *ros1* and *rdd* mutants. In contrast, 21 genes were downregulated with hypermethylated, while hypomethylated regions associated with only 4 upregulated and no downregulated genes (Fig. 3E). This asymmetry reflects the demethylase function of ROS1, as loss of its activity primarily causes methylation gains rather than losses. In addition, these expression-methylation correlations were strongly mainly CG DNA methylation-dependent manner (Fig. S6).

Among the 58 genes showing both differential expression and hypermethylation, *PIN2* was the gene directly relevant to As tolerance. *PIN2* encodes an auxin transporter previously implicated in As efflux from root cells (*25, 49*). *PIN2* transcript levels were elevated in *ros1* mutants regardless of As treatment, but increased further under As exposure (Fig. 4A). Quantitative RT-PCR confirmed this upregulation: As-treated *ros1* and *rdd* plants showed 5.09-fold and 6.99-fold increases in *PIN2* expression compared to wild type, respectively (Fig. S7A). Methylome analysis revealed that hypermethylation in *ros1* was restricted to the last exon of *PIN2*, mainly in the 3′ UTR. This region showed increased methylation— predominantly CG, with CHG and CHH also detected—while upstream regions including the promoter, 5′ UTR, and coding sequence remained unmethylated (Fig. 4A and Fig. S7B). To verify that elevated *PIN2* expression contributes to As resistance, we overexpressed *PIN2* in both wild-type and *pin2* mutant backgrounds. Both lines exhibited enhanced As tolerance (fig. S8), supporting a direct role for PIN2 in As resistance of *ros1* mutant.

**Fig. 4.**
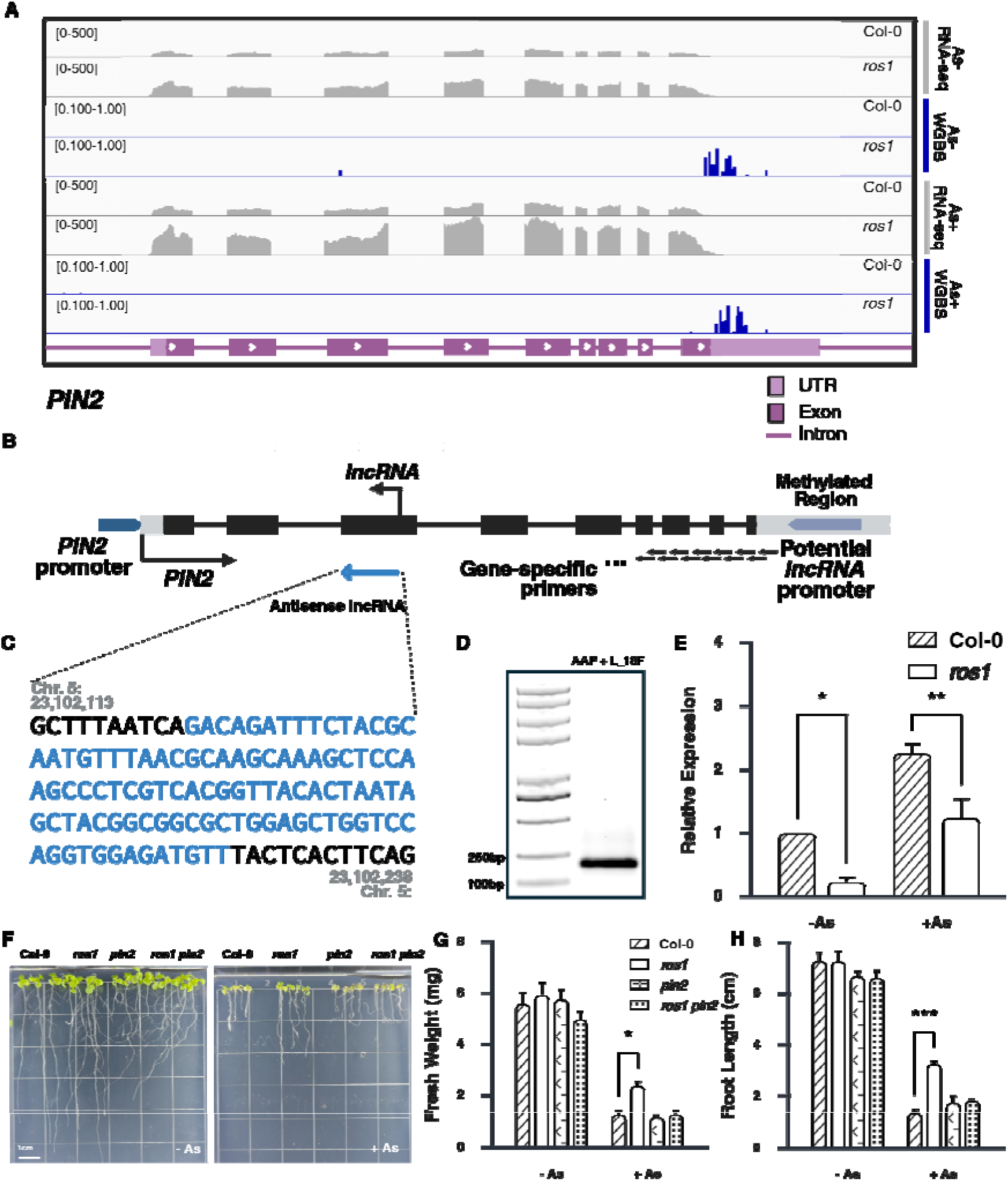
Methylation and transcription dynamics at the *PIN2* locus through an antisense lncRNA under As stress. (**A**) Genome browser view showing DNA methylation (blue) and transcript levels (gray) at the *PIN2* gene in wild-type and *ros1* mutant plants grown without or with As. (**B**) Structure of the *PIN2* locus showing the sense transcript and the antisense lncRNA mapped by 5′ RACE. (**C**) Nucleotide sequence of the antisense lncRNA amplified in (B), determined by Sanger sequencing (blue). (**D**) Gel image of the 5′ RACE product for the antisense lncRNA. Left: molecular weight marker; right: PCR product obtained with L_18F and AAP primers. (**E**) Relative expression of the antisense lncRNA in wild-type and *ros1* without or with As. (**F**) Phenotypes of wild-type, *ros1*, *pin2*, and *ros1 pin2* seedlings without (left) or with As (right). Scale bar = 1 cm. (**G**–**H**) Measurements of fresh weight (**G**) and root length (**H**). Two-way ANOVA was used for statistical comparisons. \*\*\**P* < 0.001, \*\**P* < 0.01, \**P* < 0.05.

PIN2 localizes to the plasma membrane of lateral root cap, epidermal, and cortical cells, where it exports As from the cytoplasm to the apoplast (*25, 49*). Consistent with this, *pin2* mutant showed hypersensitive to As (fig. S8). We therefore hypothesized that the As tolerance of *ros1* mutants stems from elevated *PIN2* expression caused by failed demethylation at the *PIN2* 3′ UTR. If ROS1 acts upstream of PIN2 in a linear pathway, we generated *ros1 pin2* double mutants and assessed their response to As stress. The double mutant behaved identically to *pin2* single mutants—the enhanced As tolerance of *ros1* was completely suppressed (Fig. 4F, 4G, 4H). This genetic interaction demonstrates that *PIN2* is the primary downstream effector through which *ROS1* influences As detoxification.

## An antisense lncRNA at the *PIN2* locus responds to ROS1-controlled demethylation

DNA methylation can regulate gene expression in different ways depending on where it occurs in the genome (*50, 51*). When CG methylation increases in promoters, it usually turns genes off by preventing transcription factors from binding and by attracting proteins that shut down gene activity. In contrast, CG methylation within gene bodies typically helps maintain steady gene expression and is often found in moderately active genes (*54–56*). At the *PIN2* locus, we observed higher CG methylation in the 3′ UTR in *ros1* mutants, which correlated with increased *PIN2* expression (Fig. 4A). This raised an intriguing possibility: rather than directly controlling *PIN2* transcription, this methylated region might serve as a promoter for a noncoding RNA transcribed in the opposite direction from *PIN2*.

To test this hypothesis, we employed a targeted tiling strategy using a series of forward primers spanning the *PIN2* locus, each paired with the abridged anchor primer (AAP) and abridged universal amplification primer (AUAP) in a 5′ rapid amplification of cDNA ends (5′ RACE) assay. This approach allows selective amplification of antisense transcripts while excluding sense-strand *PIN2* mRNA, as the gene specific primers would only generate products from templates transcribed in the opposite direction. By systematically tiling primers across the locus, we aimed to map the transcription start site of any putative antisense RNA and determine whether its origin coincides with the hypermethylated region in *ros1* mutants (Fig. 4B). This approach revealed an antisense long noncoding RNA (lncRNA) that overlaps with *PIN2*’s third exon, which we verified through Sanger sequencing (Fig. 4A-C).

Antisense lncRNAs are increasingly recognized as regulatory molecules that can silence their corresponding sense transcripts through direct RNA–RNA pairing, triggering degradation or blocking translation (*57–59*). Based on this precedent, we propose that methylation at the *PIN2* 3′ UTR controls expression of the antisense lncRNA, which regulates *PIN2* mRNA levels. To confirm this regulatory relationship, we measured antisense lncRNA levels in wild-type and *ros1* plants using quantitative RT-PCR with 5’-RACE products as template. and observed that its expression pattern was inversely correlated with *PIN2* mRNA abundance (Fig 4E). This opposite expression pattern between the antisense lncRNA and *PIN2* mRNA strongly suggests that the antisense RNA directly controls *PIN2* expression levels.

## Sense and antisense transcription at *PIN2* are regulated through distinct chromatin mechanisms

To confirm that ROS1-mediated demethylation controls *PIN2* expression by antisense lncRNA transcription, we also examined the chromatin landscape at this locus using ChIP-seq datasets. We first asked where ROS1 binds within the *PIN2* gene in wild-type plants. Myc-ROS1 ChIP-seq data revealed ROS1 occupancy throughout the entire *PIN2* locus, including the last exon of PIN2 region where we observed hypermethylation in *ros1* (Fig. 5A) (*60*).

**Fig. 5.**
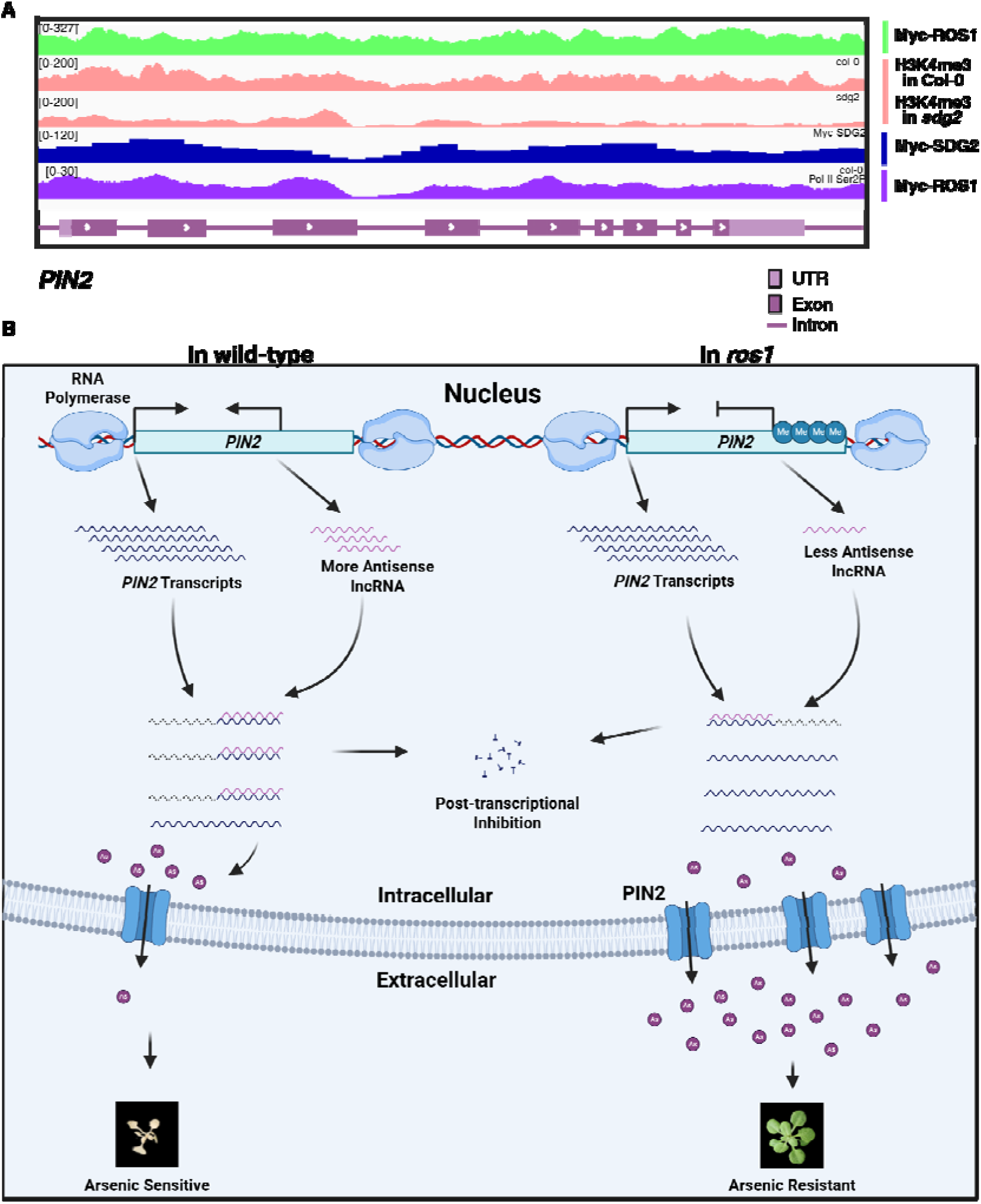
Chromatin landscape and antisense lncRNA at the *PIN2* locus under As stress. (**A**) ChIP-seq analysis of the *PIN2* genomic region showing enrichment patterns for Myc- ROS1, H3K4me3, Myc-SDG2, and RNA polymerase II. (**B**) Model of *ROS1*-dependent regulation of *PIN2* under As stress.

Since H3K4me3 serves as the master recruiting signal for ROS1-mediated DNA demethylation (*60*), we next examined H3K4me3 distribution at *PIN2*. H3K4me3 marks were detected throughout the *PIN2* gene body, including the third exon where the antisense lncRNA is expressed. The *PIN2* promoter showed H3K4me3 enrichment extending from the transcription start region through the gene body while remaining unmethylated—consistent with active gene promoters (Fig. 5A). These regions also displayed RNA polymerase II occupancy and Myc-ROS1 binding, confirming active *PIN2* transcription and demethylase recruitment (*61*). We also examined the distribution of SET DOMAIN GROUP2 (SDG2), the major H3K4 methyltransferase in Arabidopsis. Myc-SDG2 ChIP-seq data showed SDG2 occupancy across all *PIN2* regions, paralleling the H3K4me3 distribution (*62*). To determine whether H3K4me3 deposition at the antisense lncRNA start site depends on SDG2, we analyzed ChIP-seq data from *sdg2* mutants. Loss of *SDG2* caused widespread reduction in H3K4me3 levels throughout most of the *PIN2* gene body (*60*). Interestingly, however, a distinct H3K4me3 peak persisted at the third exon where the antisense lncRNA transcription initiates (Fig. 5A). This residual H3K4me3 peak observed mainly at the third exon, indicating the site-specific H3K4me3 in antisense lncRNA region is independent of SDG2 and suggesting that *PIN2* sense and antisense lncRNA transcription are regulated through distinct mechanisms.

Together, these chromatin profiling data define a regulatory landscape at *PIN2* where ROS1- mediated demethylation, guided by H3K4me3 marking, maintains low methylation levels throughout the gene body in wild-type plants. At the 3′ UTR, this demethylation activity specifically suppresses antisense lncRNA transcription. When ROS1 is absent, methylation accumulates at this region, silencing the antisense lncRNA and thereby releasing *PIN2* sense transcription from negative regulation (Fig. 5B). This model explains how methylation changes at the *PIN2* 3′ UTR indirectly control *PIN2* expression by modulating antisense lncRNA levels, thereby regulating As tolerance (Fig. 5B).

## Discussion

We demonstrate that *ROS1*-mediated DNA demethylation regulates As tolerance in Arabidopsis through epigenetic control of *PIN2*. Loss of *ROS1* causes hypermethylation at the *PIN2* 3′ UTR, which suppresses an antisense lncRNA and consequently increases *PIN2* expression. Transcriptomic analysis revealed that relatively few genes showed differential expression in *ros1* mutants, suggesting that *ROS1* regulates As tolerance through specific rather than genome-wide changes. Notably, expression patterns in *ros1* and the triple mutant *rdd* were highly similar following As treatment, whereas *dml2* and *dml3* single mutants showed different profiles. This similarity indicates that *ROS1* plays the primary role in As tolerance, while *DML2* and *DML3* contribute less. GO and GSEA analysis showed that *dml2* and *dml3* are more involved in plant vegetative and reproductive development (Fig. 2E and fig. S2 and fig. S3 and table S2), suggesting functional specialization among DNA demethylases that may enable flexible epigenetic responses.

Through integrated transcriptomic and methylome analyses, we identified *PIN2* as a direct target of *ROS1* during As stress, a finding we confirmed genetically using *pin2 ros1* double mutants. *PIN2* encodes an auxin efflux carrier that mediates polar auxin transport and controls root gravitropism, and also exports As from root cells (*63, 64*). Notably, hypermethylation at the *PIN2* 3′ UTR in *ros1* mutants occurred even without As treatment and correlated with elevated *PIN2* transcript levels. In wild-type plants, lower *PIN2* expression corresponded with higher antisense lncRNA transcription from the same genomic region. In *ros1* mutants, hypermethylation suppressed antisense lncRNA production, thereby derepressing *PIN2*. This regulatory pathway potentially connects *ROS1*-mediated demethylation to auxin distribution and root architecture through *PIN2* control, indicating that future work should examine how DNA demethylation influences auxin homeostasis via *PIN2* regulation.

Natural antisense lncRNAs, transcribed from the opposite strand of their associated genes, regulate expression through multiple mechanisms. These transcripts respond to epigenetic modifications including DNA methylation and histone marks, and can alter chromatin structure, RNA stability, splicing, or translation (*65*). Our model proposes that the *PIN2* antisense lncRNA represses *PIN2* expression, possibly through double-stranded RNA formation. Without *ROS1*, hypermethylation accumulates at the *PIN2* 3′ UTR, which functions as the antisense lncRNA promoter. This methylation reduces antisense lncRNA production, releasing *PIN2* from repression and increasing its transcript levels above wild type. Enhanced *PIN2* protein at the plasma membrane improves As efflux, conferring tolerance in *ros1* mutants (Fig. 5B). This regulatory strategy—where gene body methylation at the 3′ UTR indirectly activates gene expression by silencing an inhibitory antisense lncRNA—appears common among natural antisense transcripts (*66, 67*).

The retention of H3K4me3 marks at the antisense transcription start site in *sdg2* mutants offers critical evidence. This observation indicates that the antisense lncRNA represents a genuinely regulated transcript rather than random transcriptional noise, since non-functional transcription typically lacks defined chromatin modifications (*68*). The antisense start site shows a focused H3K4me3 peak typical of authentic promoters, even when the major H3K4 methyltransferase SDG2 is absent. This SDG2-independent marking suggests alternative H3K4 methyltransferases or cooperative with SDG2 establish H3K4me3 at this site, allowing antisense transcription to occur through distinct chromatin mechanisms compared to sense *PIN2* transcription (*69*). Such regulatory independence may allow the antisense lncRNA to respond to environmental signals in ways that are distinct from the sense transcript, enabling flexible control of *PIN2* expression. Together, our findings reveal a novel epigenetic pathway linking DNA demethylation to heavy metal tolerance and suggest potential strategies for engineering stress-resilient plants on contaminated soils.

## Acknowledgments

This study was supported by the Startup fund at Duke Kunshan University, 2023, 2024, 2025 Summer Research Scholarship fund, 2024, 2025 Summer Research Grant, 2021 Interdisciplinary Seed Grant, Synear and Wang-Cai Seed Grant 2022, and 2022, 2023, 2024, 2025 Wang-Cai Biochemistry Lab Grant, 2025 Duke University Provost fund.

## Author contributions

J.L., L.S., Y.W., and Y. Yang conceived the project and designed the research; J.L. and L.S. acquired the funding for this work; X.C., Y.W., Y. Yang, and Z.H. performed the original phenotypical assays; X.L., Y.W., Z.H., and Z.Y prepared the WGBS and whole transcriptome sequencing; J.Y., Y.W., Z.H., and Z.Y. analyzed the next generation sequencing data; R.W., Y.W., Y. Yao, and Z.H. constructed the double mutant and overexpression lines; R.W., Y.W., Y. Yao, Z.X and Z.H conducted the 5’ RACE and Sanger sequencing assays; H.Y., Y.W., and Z.X. designed and performed quantitative PCR assays; Y.W. wrote the original draft; H.Y., J.L. L.S., R.W., Y.W., Y. Yao, Z.H., and Z.X. revised the manuscript; J.L. and L.S. supervised the project.

## Data and materials availability

All data are available in the main text or the supplementary materials. The WGBS and whole transcriptome sequencing raw data have been deposited in the National Center for Biotechnology Information Gene Expression Omnibus (accession #).

## Materials and Methods

### Plant material and growth conditions

*Arabidopsis thaliana* Columbia-0 (Col-0) was used as the wild-type control throughout this study. We used previously characterized DNA demethylase mutants including *ros1-4*, *dml2-2*, *dml3-2*, and the triple mutant *rdd-2* (70), as well as the auxin transport mutant *pin2* (SALK_122916C) (*71*). To create the *ros1 pin2* double mutant, we performed crosses between *ros1-4* and *pin2* plants and confirmed homozygous double mutants by genotyping PCR. All oligonucleotide primers used in this study are listed in Table S1.

Seeds were surface-sterilized by treatment with 50% (v/v) bleach solution and washed three times with sterile distilled water. Seeds were then plated on half-strength Murashige and Skoog (1/2 MS) medium solidified with agar (Coolaber, China) and cold-treated at 4°C for 2 days to promote uniform germination. For phenotypic characterization, we transferred 5-day-old seedlings to 1/2 MS medium supplemented with 200 μM or 250 μM Na HAsO for As stress treatment. For genome-wide transcriptome and methylome analyses, 14-day-old seedlings were transferred to 1/2 MS medium containing 250 μM Na HAsO and grown for 7 more days before harvest. All plates were oriented vertically in a growth chamber (AR1200, Wuhan Ruihua, China) set to 22°C with 16-hour light/8-hour dark cycles.

### Transgenic plant generation and growth measurements

To create complementation lines, the complete coding sequences of *ROS1* or *PIN2* were amplified and inserted into the Gateway entry vector pDONR221 (Invitrogen, USA), then transferred into the expression vector pGWB502 through LR recombination (*74*). These constructs were introduced into *Agrobacterium tumefaciens* strain GV3101 and used to transform *ros1-4*, *pin2*, or Col-0 plants following established floral dip procedures (*75*).

We measured primary root length from digital photographs using ImageJ software (version 1.54). Fresh weight was measured by removing seedlings from culture medium, gently removing surface moisture with tissue paper, and weighing on a Sartorius Quintix analytical balance (Sartorius AG, Germany). Each experiment included four biological replicates, and statistical analysis was performed using two-way ANOVA.

### Gene expression analysis by quantitative PCR

Total RNA was purified from plant tissues with the FastPure Plant Total RNA Isolation Kit (Vazyme Biotech, China) according to the manufacturer’s protocol. We assessed RNA concentration and purity using a NanoDrop 2000 spectrophotometer (Thermo Fisher, USA). Complementary DNA was synthesized from 5 ng total RNA using the HiScript II 1st Strand cDNA Synthesis Kit (Vazyme Biotech, China) with both oligo(dT) and random hexamer primers (Qiagen, Germany).

Expression levels were quantified by SYBR Green-based qPCR using the SYBR Green/ROX qPCR Master Mix (Vazyme Biotech, China) with cDNA from arsenic-treated and control samples. Only reactions with a single melting peak between 75°C and 85°C were accepted for analysis. For the antisense *PIN2* lncRNA, we performed probe-based qPCR using Taq Pro U+ multiple probe qPCR mix (Vazyme Biotech, China) with 5′ RACE-amplified products as templates. Sequences for all primers and probes are provided in Table S1. Expression values were normalized to the reference gene *CBP20*. Amplifications were run on a QuantStudio 6 Pro Real-Time PCR System (Applied Biosystems, USA), and relative expression was determined by the 2^^−ΔΔCt^ method (*76*).

### 5′ RACE assay and tiling PCR

We designed a panel of primers covering the *PIN2* genomic region (positions 23,101,000– 23,104,500) with an average spacing of 200 bp (Table S1). In regions corresponding to the third exon and 3′ UTR of *PIN2*, primer spacing was reduced to 100 bp for higher resolution mapping. We performed 5′ RACE using 5 ng total RNA (Qiagen, Hilden, Germany).

Following reverse transcription and RNase H digestion, cDNA molecules were extended with poly-C tails using terminal deoxynucleotidyl transferase (Sangon Biotech, China). Nested PCR was conducted using gene-specific primers combined with an abridged anchor primer (AAP) and abridged universal amplification primer (AUAP), with amplification performed using high-fidelity KOD DNA polymerase (Toyobo, Osaka, Japan). PCR products were separated on 1.5% agarose gels, purified, and analyzed by Sanger sequencing.

### RNA sequencing and computational analysis

Twenty-one-day-old seedlings (14 days under control conditions plus 7 days with or without 250 μM arsenic treatment) were collected and snap-frozen in liquid nitrogen. Total RNA was extracted using TRIzol reagent (Invitrogen, Thermo Fisher Scientific, Waltham, MA, USA), and cDNA libraries were prepared with the TruSeq RNA Sample Preparation Kit (Illumina, San Diego, CA). Two biological replicates were generated per condition. Libraries were sequenced in paired-end mode on an Illumina HiSeq 2000 platform (Illumina, San Diego, CA).

Adapter removal and quality trimming were performed using Cutadapt v2.8, and read quality was verified with FastQC v0.11.9 (*77, 78*). Trimmed reads were aligned to the *Arabidopsis* TAIR10 reference genome using HISAT2 (*79*). Differential expression analysis was conducted with DESeq2 in R, with genes showing absolute fold change ≥ 2 and adjusted *P*- value ≤ 0.05 considered significantly differentially expressed (*80*). For transposable element analysis, reads were mapped to the Araport11 transposable element annotation, and differential expression was assessed using DESeq2 with the same statistical cutoffs.

### Bisulfite sequencing and DNA methylation analysis

We extracted genomic DNA from plant samples and fragmented it by sonication. Fragmented DNA was end-repaired and A-tailed to prepare for adapter ligation. Unmethylated lambda DNA was added to each sample as a spike-in control for assessing bisulfite conversion efficiency. Samples were treated with sodium bisulfite to convert unmethylated cytosines to uracil while preserving methylated cytosines. Converted DNA was amplified and sequenced on an Illumina HiSeq 2000 platform (Illumina, San Diego, CA).

Raw reads were processed with Cutadapt v2.8 to remove adapters and low-quality bases, and quality was checked using FastQC v0.11.9 (*77, 78*). We aligned bisulfite-treated reads to a converted version of the *Arabidopsis* TAIR10 reference genome using Batmeth2 (*81*). We aligned bisulfite-treated reads to a converted version of the *Arabidopsis* TAIR10 reference genome using Batmeth2 (*81*). Differentially methylated genes were identified with methylKit in R (*82*). We applied significance thresholds of 40% methylation difference for CG sites, 20% for CHG sites, and 10% for CHH sites, all with adjusted *P*-value ≤ 0.05.

---

**Fig. S1.**
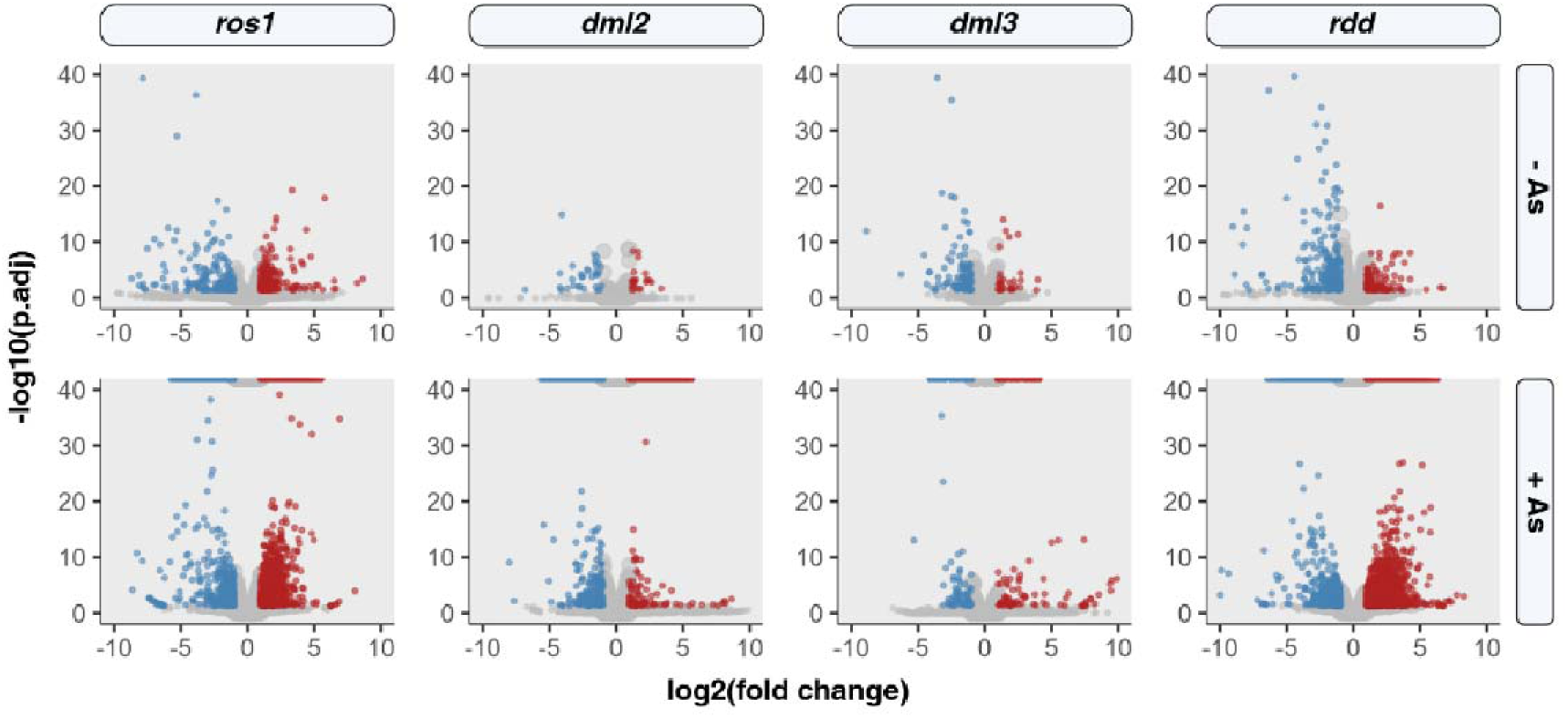
Gene expression changes in demethylase mutants with and without As treatment. Top row shows control conditions (-As); bottom row shows arsenic-treated samples (+As) for *ros1*, *dml2*, *dml3*, and *rdd*. Red dots denote upregulated genes; blue dots denote downregulated genes (|log FC| > 2, adjusted *p* < 0.05).

**Fig. S2.**
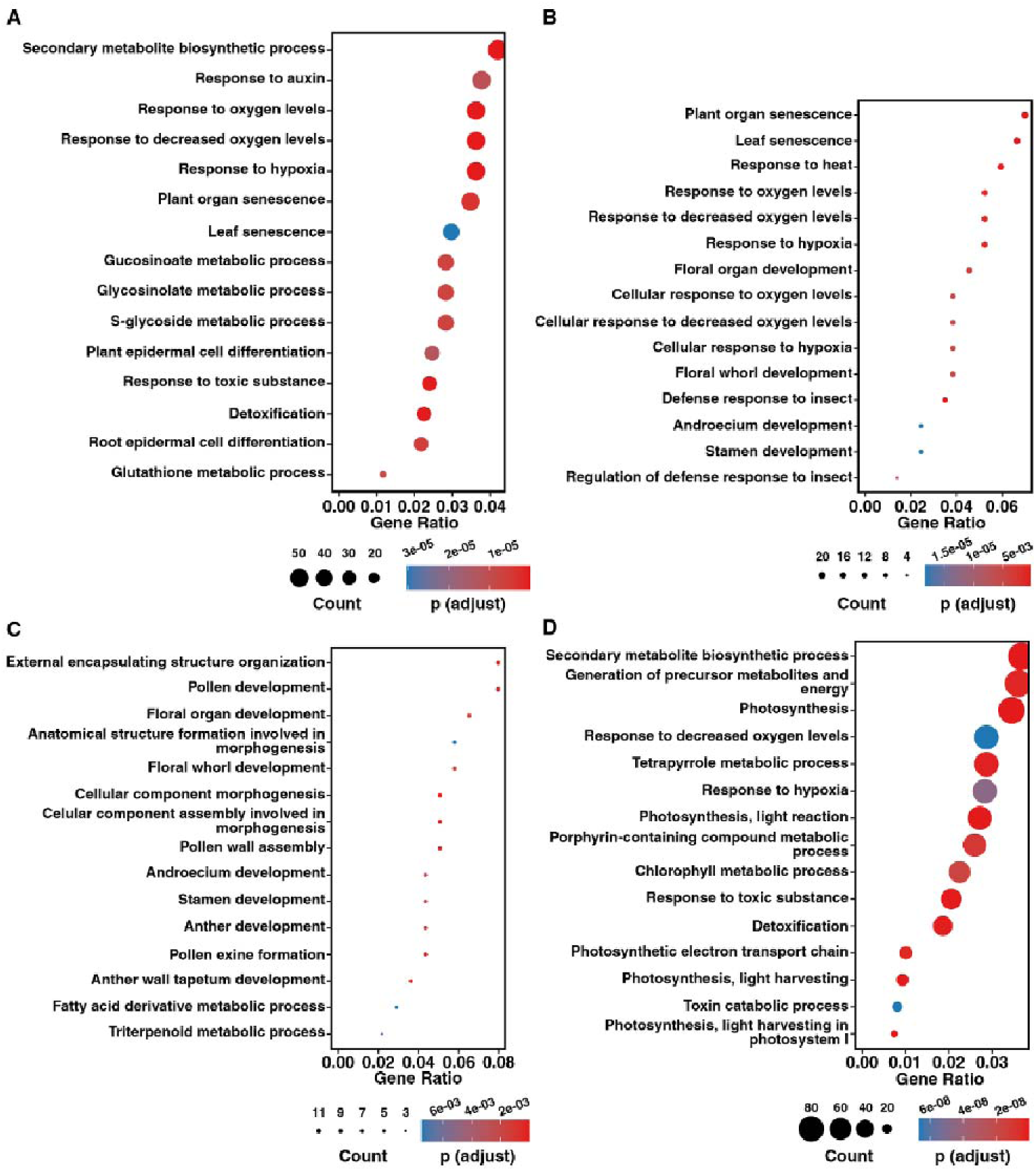
Functional enrichment of differentially expressed genes under As stress. Bubble plots present significantly enriched gene ontology terms for each mutant background: *ros1* (**A**), *dml2* (**B**), *dml3* (**C**), and *rdd* (**D**).

**Fig. S3.**
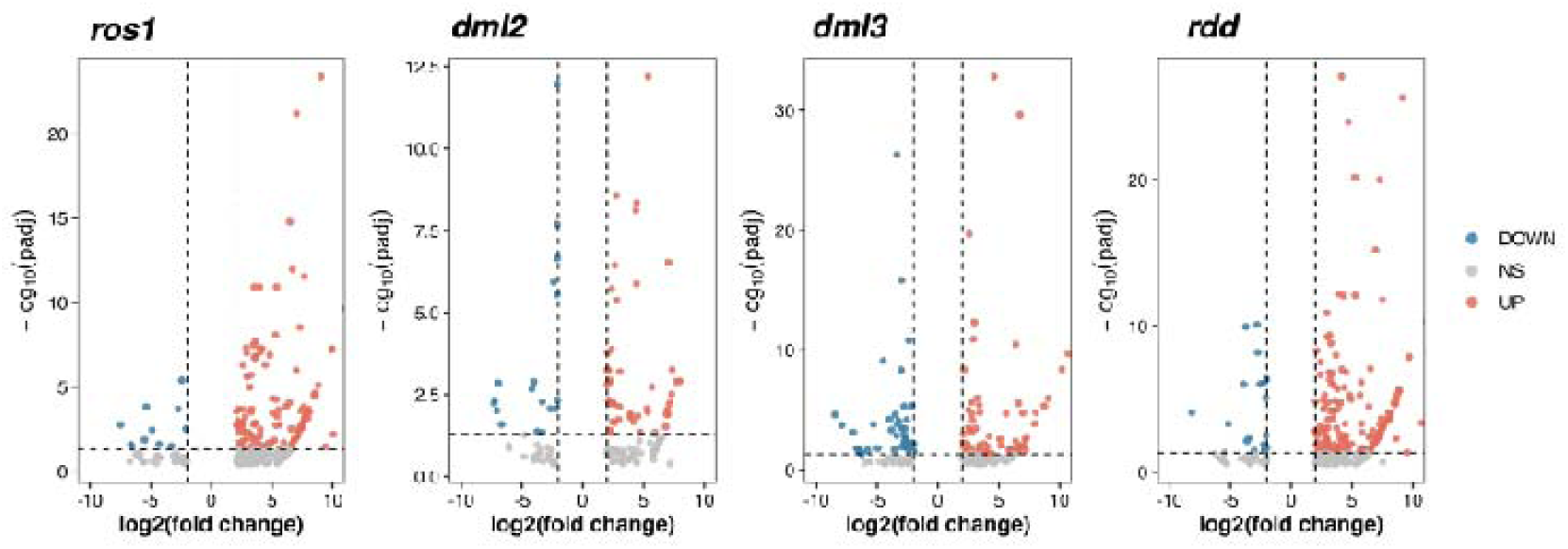
Differential transposon expression in *ros1*, *dml2*, *dml3*, and *rdd* under As stress. Expression changes are shown for *ros1*, *dml2*, *dml3*, and *rdd* mutants, with red and blue dots representing upregulated and downregulated transposons, respectively (|log FC| > 2, adjusted *p* < 0.05).

**Fig. S4.**
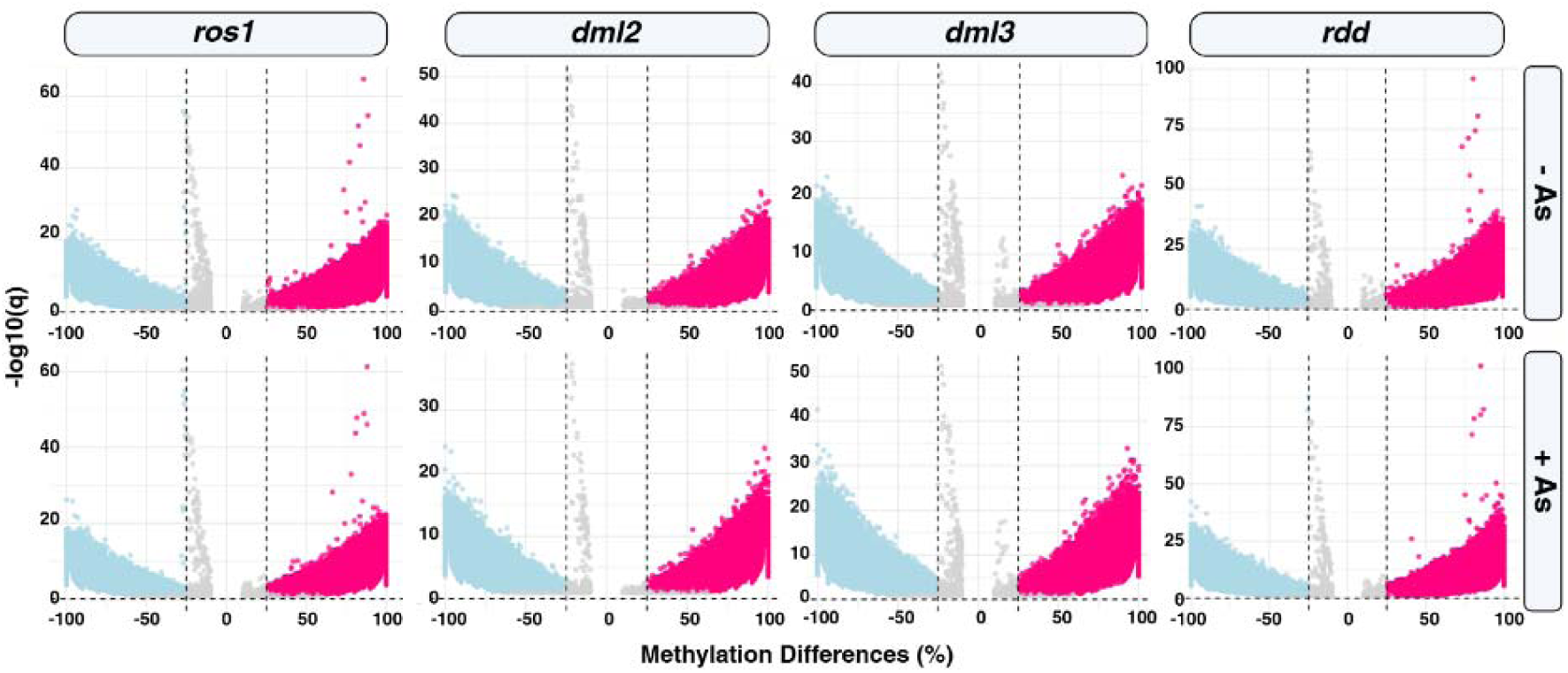
Genome-wide methylation alterations in demethylase mutants with and without As treatment. Hypomethylated genes (blue dots) and hypermethylated genes (red dots) are displayed for control (-As, top row) and arsenic-treated (+As, bottom row) conditions. Methylation difference threshold = 25%.

**Fig. S5.**
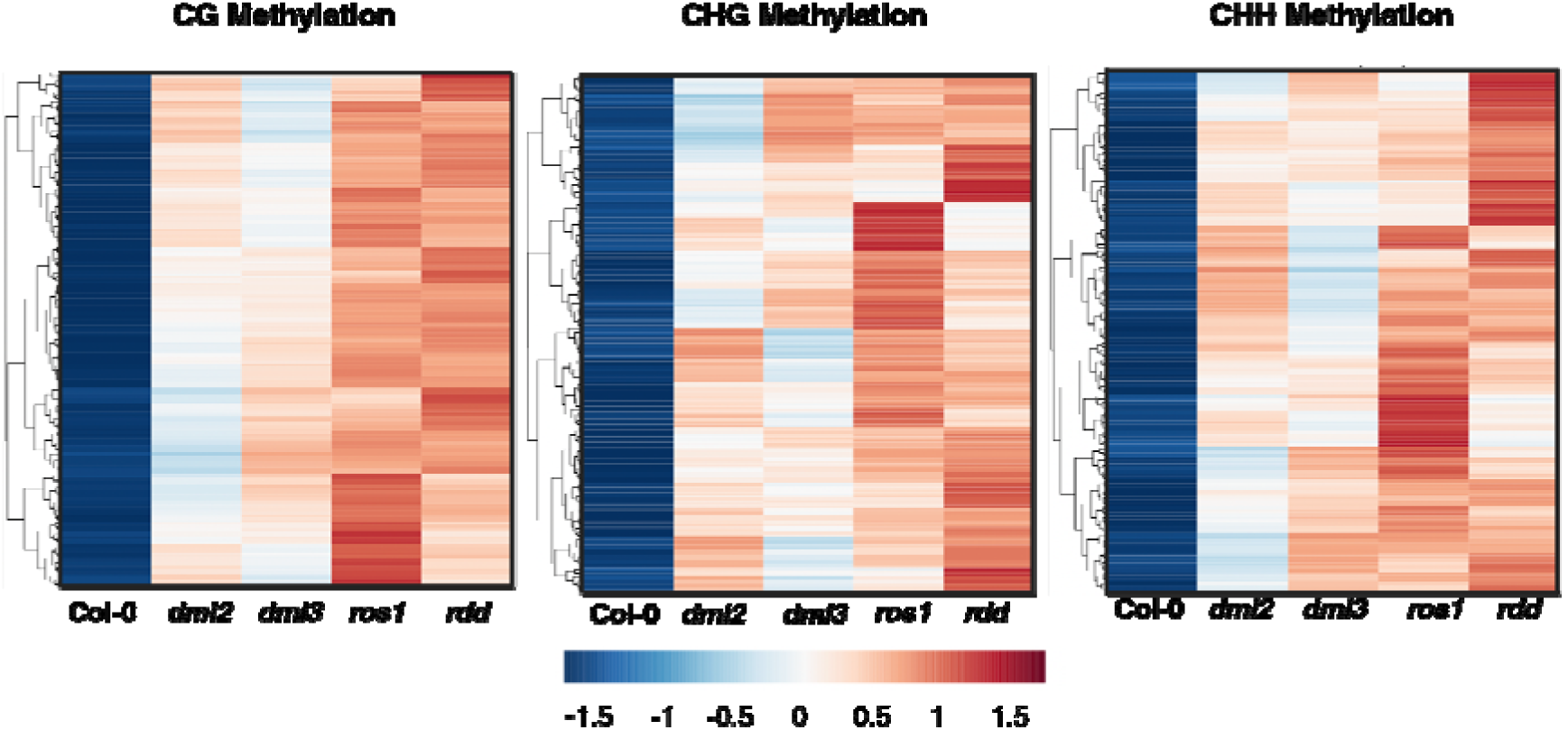
DNA methylation patterns across wild-type, *dml2*, *dml3*, *ros1*, and *rdd* under As stress. CG, CHG, and CHH methylation levels are compared across wild-type, *dml2*, *dml3*, *ros1*, and *rdd* genotypes.

**Fig. S6.**
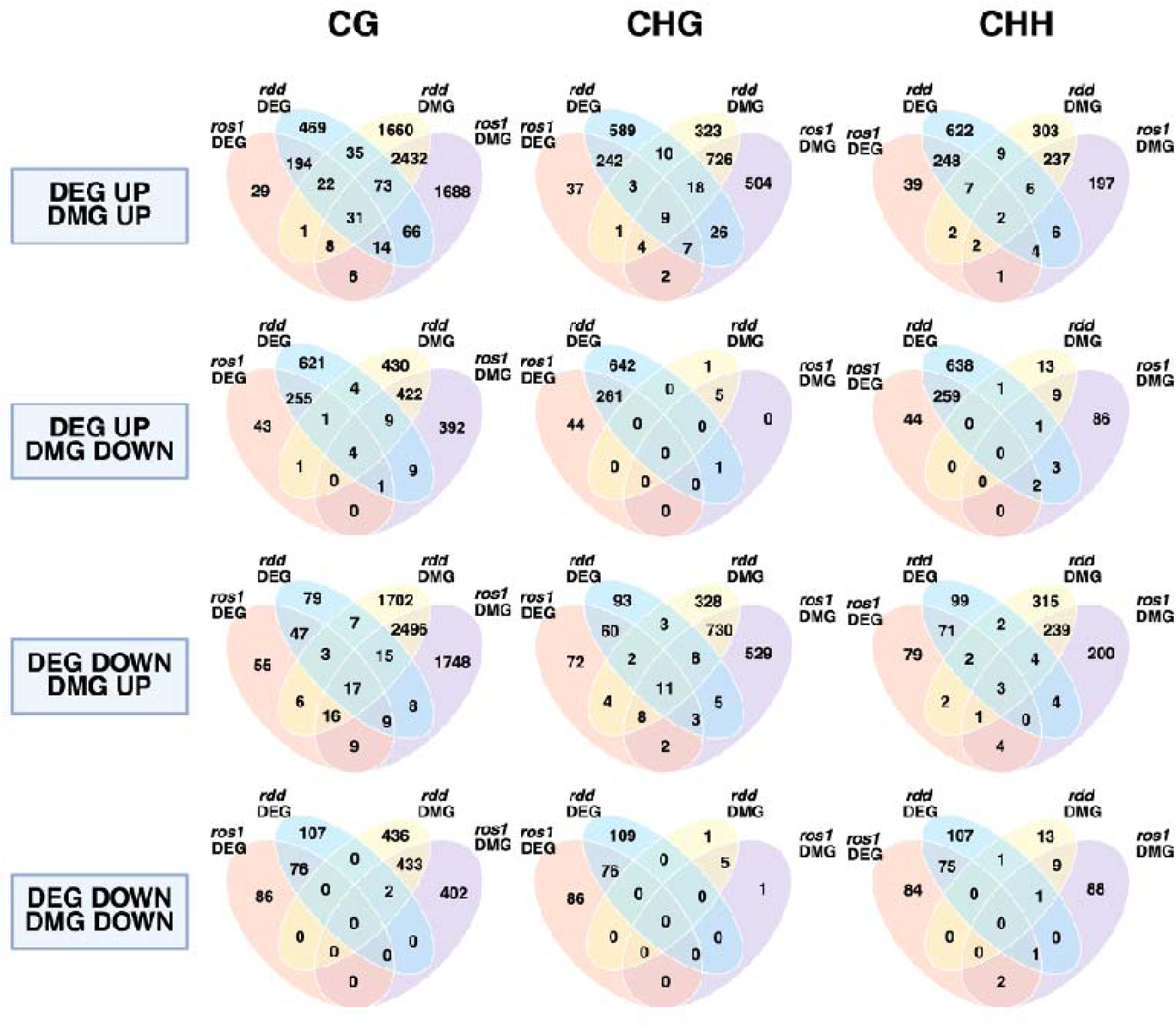
Integration of transcriptome and methylome data between *ros1* and *rdd* under As stress. Overlapping genes showing both differential expression and differential methylation in *ros1* and *rdd* mutants are presented for each methylation context (CG, CHG, CHH).

**Fig. S7.**
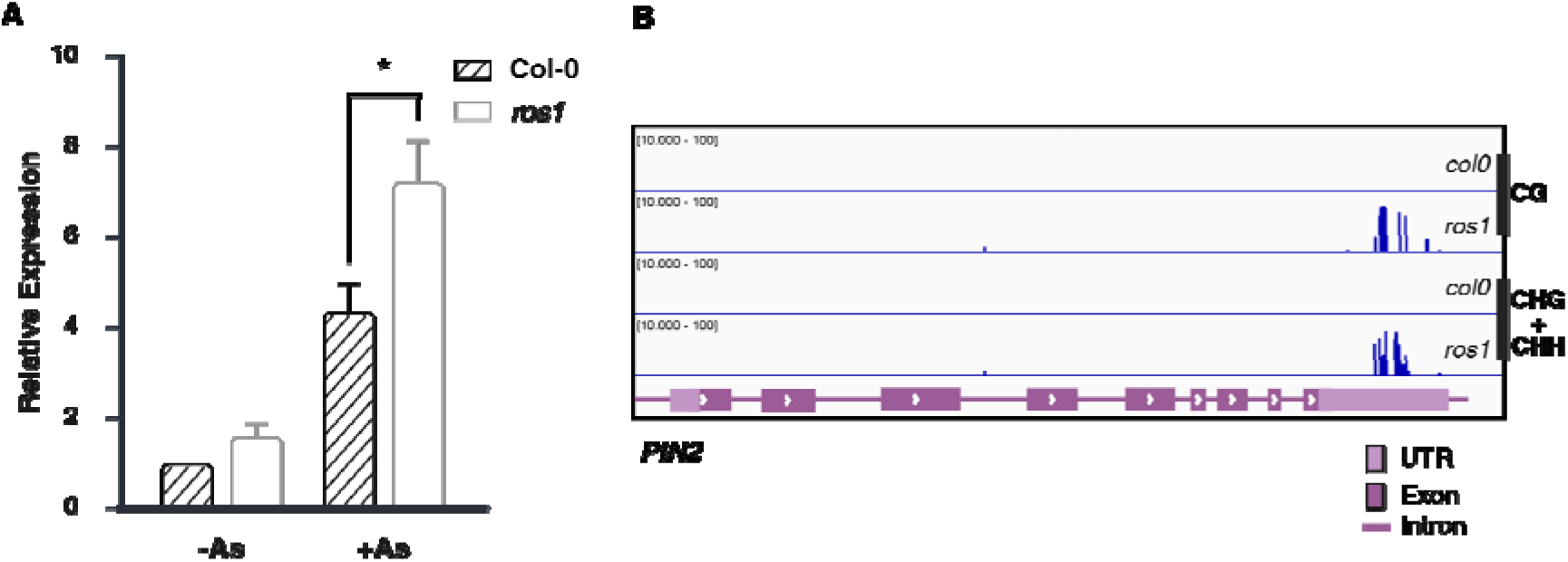
ROS1-dependent regulation of *PIN2*. (**A**) qRT-PCR analysis of *PIN2* transcript levels in wild-type and *ros1* with or without As treatment. (**B**) Integrative Genomics Viewer tracks displaying methylation levels at the *PIN2* locus in both CG and combined CHG+CHH contexts.

**Fig. S8.**
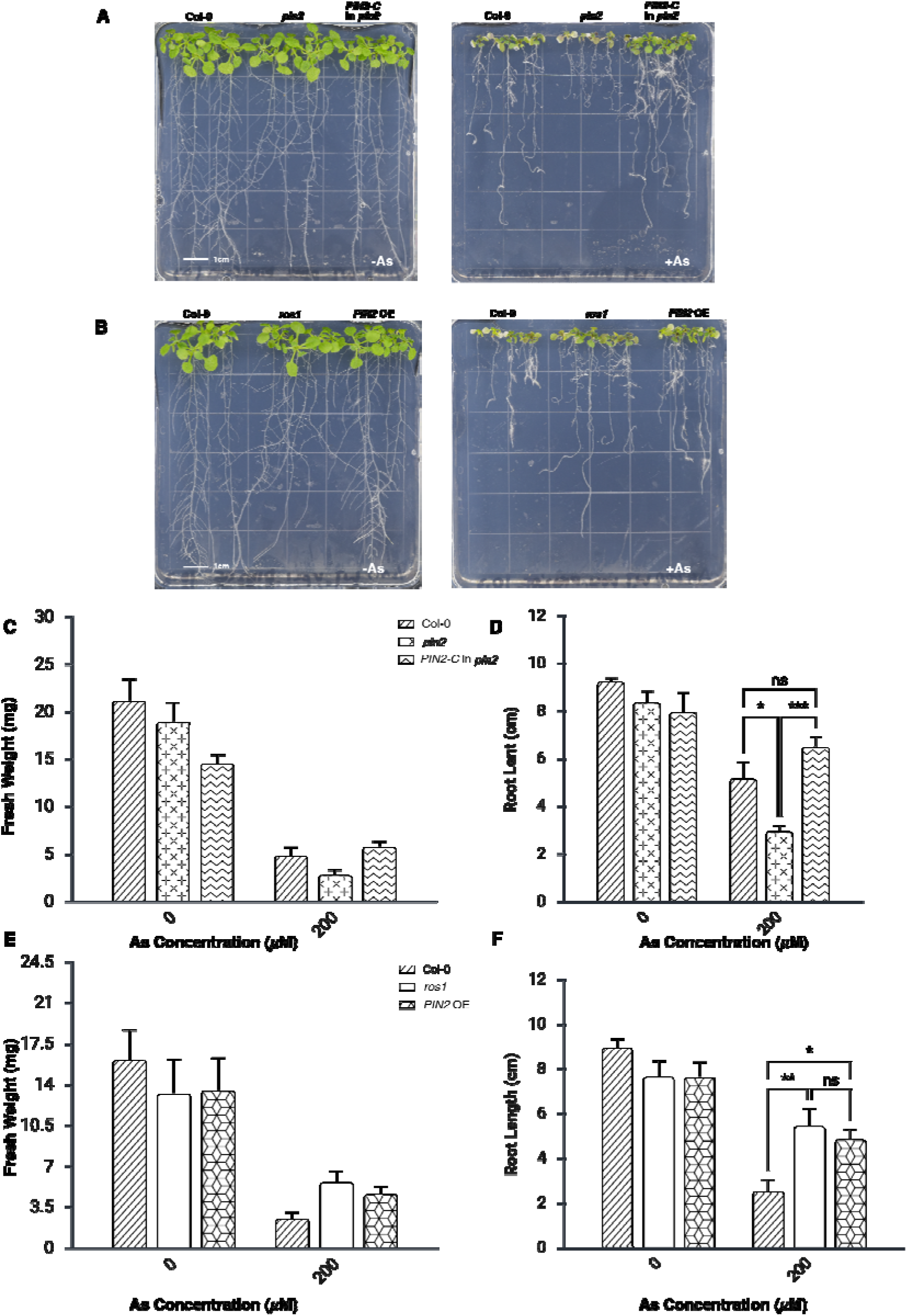
Genetic complementation and overexpression phenotypes under As stress. (**A**) Growth phenotypes of wild-type (Col-0), *pin2* mutant, and complemented *pin2* (*PIN2*-C in *pin2*) line without or with As exposure. (**B**) Growth phenotypes of wild-type, *ros1* mutant, and *PIN2* overexpression line (*PIN2* OE) without or with As exposure. Scale bars = 1 cm. (**C**–**D**) Quantification of fresh weight (**C**) and primary root length (**D**). (**E**–**F**) Quantification of fresh weight (**E**) and primary root length (**F**). Statistical significance was assessed by two-way ANOVA. *\**, *P* < 0.001; **, *P* < 0.01; *, *P* < 0.05.

**Table S1. Primers and probes used in this study.**

**Table S2. GSEA enrichment terms of DEGs in *ros1*, *dml2*, *dml3*, and *rdd*.**

